# Perceptual uncertainty alternates top-down and bottom-up fronto-temporal network signaling during response inhibition

**DOI:** 10.1101/2021.12.27.474302

**Authors:** Kaho Tsumura, Reiko Shintaki, Masaki Takeda, Junichi Chikazoe, Kiyoshi Nakahara, Koji Jimura

**Affiliations:** Department of Biosciences and Informatics, Keio University, Yokohama, Japan; Research Center for Brain Communication, Kochi University of Technology, Kami Japan; Supportive Center for Brain Research, National Institute for Physiological Sciences, Okazaki, Japan

## Abstract

Response inhibition is a primary executive control function that allows the withholding of inappropriate responses, and requires appropriate perception of the external environment to achieve a behavioral goal. It remains unclear, however, how response inhibition is achieved when goal-relevant information involves perceptual uncertainty. Twenty-six human participants of both sexes performed a go/no-go task where visually presented random-dot motion stimuli involved perceptual uncertainties. The right inferior frontal cortex (rIFC) was involved in response inhibition, and the middle temporal (MT) region showed greater activity when dot motions involved less uncertainty. A neocortical temporal region in the superior temporal sulcus (STS) specifically showed greater activity during response inhibition in more perceptually certain trials. In this STS region, activity was greater when response inhibition was successful than when it failed. Directional effective connectivity analysis revealed that in more coherent trials, the MT and STS regions showed enhanced connectivity to the rIFC, whereas in less coherent trials, the signal direction was reversed. These results suggest that a reversible fronto-temporal functional network guides response inhibition under perceptual uncertainty, and in this network, perceptual information in the MT is converted to control information in the rIFC via STS, enabling achievement of response inhibition.

**Significance statement:** Response inhibition refers to withholding inappropriate behavior and is an important for achieving goals. Often, however, decision must be made based on limited environmental evidence. We showed that successful response inhibition is guided by a neocortical temporal region that plays a hub role in converting perceived information coded in a posterior temporal region to control information coded in the prefrontal cortex. Interestingly, when a perceived stimulus becomes more uncertain, the prefrontal cortex supplements stimulus encoding in the temporal regions. Our results highlight fronto-temporal mechanisms of response inhibition in which conversion of stimulus-control information is regulated based on the uncertainty of environmental evidence.

## Introduction

Inhibition of inappropriate responses is a representative executive control function guiding adaptation to changing environments (Miller and Cohen, 2001; Aron et al., 2004; Dalley et al., 2004; Ridderinkhof et al., 2004). Prior neuropsychological, electrophysiological, and neuroimaging studies have identified critical brain regions associated with response inhibition in the fronto-parietal cortex and subcortical nucleus (Garavan et al., 1999; Konishi et al., 1999; Bokura et al., 2001; Braver et al., 2001; Menon et al., 2001; Aron et al., 2004; Aron and Poldrack, 2006; Li et al., 2006; Aron et al., 2007; Chikazoe et al., 2007; Li et al., 2008; Xue et al., 2008; Chambers et al., 2009; Chikazoe et al., 2009; Swann et al., 2009; White et al., 2014; Osada et al., 2019). Among these regions, the right inferior frontal cortex (rIFC) is thought to be responsible for a core mechanism underlying response inhibition (Aron et al., 2004; Chikazoe et al., 2007). The aforementioned studies assumed that task-relevant information is appropriately extracted by the perception of external environments, and used behavioral tasks involving perception of distinctive sensory stimuli (Fig 1A). In our daily life, however, goal-relevant environmental information is not always available or distinctive. It is thus unclear how response inhibition is achieved when environmental information involves perceptual uncertainties.

**Figure 1.**
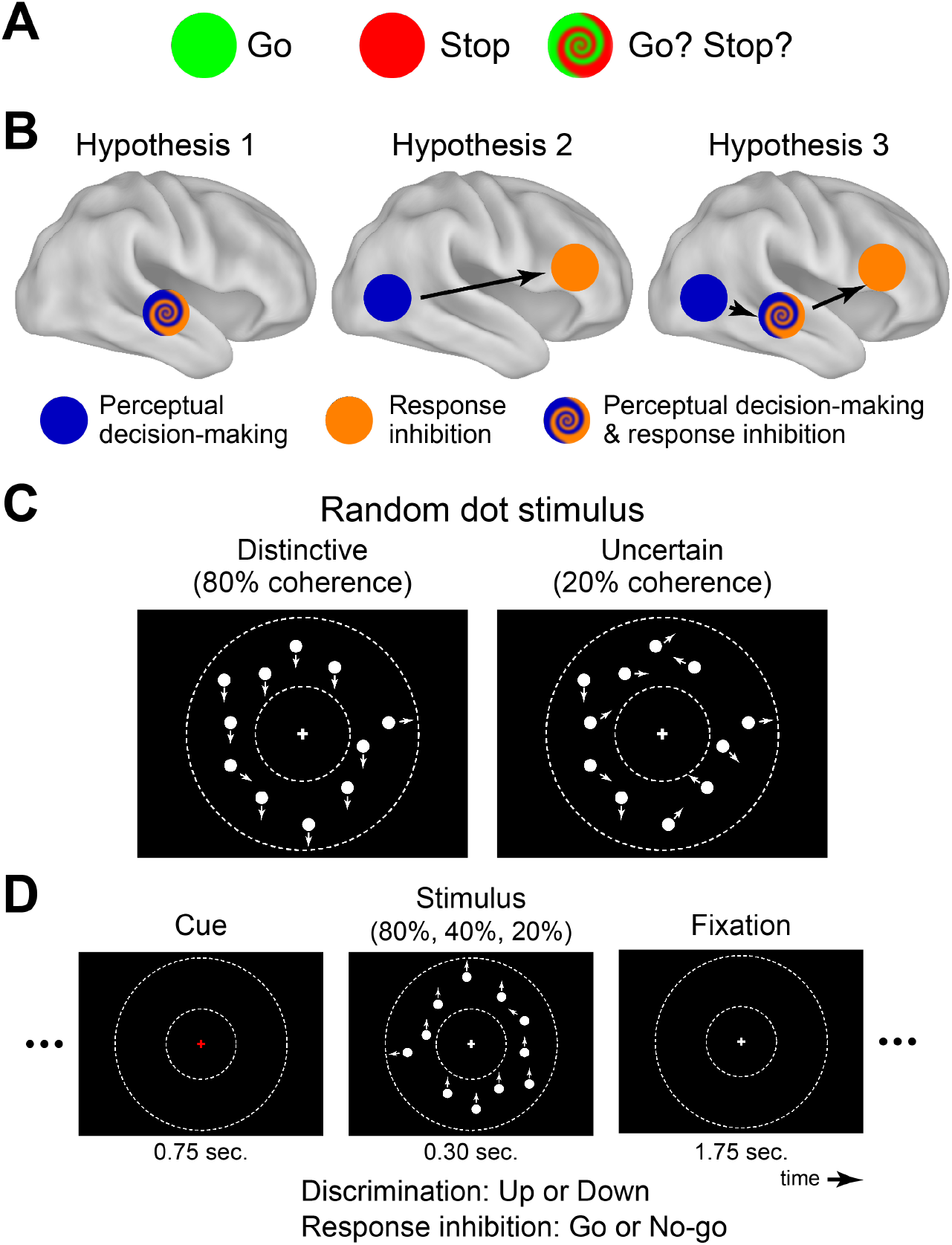
A) Response inhibition and perceptual uncertainty. B) Three possible functional topological structures of response inhibition under perceptual uncertainty. C) Random dot motion stimulus. White dots were presented within a donut-shaped display circle as indicated by dotted lines. Each arrow indicates the direction of motion of its corresponding dot. D) Discrimination task and go/no-go tasks. Motion dot stimuli were cued by a red fixation cross followed by a white fixation cross.

Perceived sensory information from the external environment guides a course of action, a process referred to as perceptual decision-making. This function involves extraction of relevant sensory information and making suitable decisions. Studies of perceptual decision-making have used behavioral tasks that demand discrimination of presented sensory stimuli (Newsome and Paré, 1988; Corbetta et al., 1991; Erwin et al., 1992; Romo et al., 1998). In a common task, randomly moving dots are visually presented and motion strength (coherence) is manipulated by the proportion of dots moving coherently toward one direction. Behavioral agents are then required to judge the overall direction of movement of the dot stimuli. It has been shown that, when a low-coherence dot stimulus is presented, accuracy is reduced and reaction times (RTs) become longer (Shadlen et al., 1996; Kim and Shadlen, 1999; Mazurek et al., 2003; Palmer et al., 2005; Kayser et al., 2010). Electrophysiological and neuroimaging studies have shown that perception of moving dots is associated with the middle temporal (MT) cortex, with greater activity when the motion coherence is high (Newsome and Paré, 1988; Britten et al., 1992, 1993; Zohary et al., 1994; Shadlen et al., 1996; Beauchamp et al., 1997; Britten and Newsome, 1998; Kim and Shadlen, 1999; Braddick et al., 2001; Corbetta and Shulman, 2002; Huk et al., 2002; Mazurek et al., 2003; Kayser et al., 2010). Importantly, MT activity is enhanced when behavioral agents actively attend to the movement (Beauchamp et al., 1997; Corbetta and Shulman, 2002; Mante et al., 2013; Zhang et al., 2013; Kumano et al., 2016), suggesting that MT involvement reflects the perception of goal-relevant information rather than simple accumulation of stimulus evidence. As such, the MT plays an important role in perceptual decision-making regarding moving stimuli.

Given the characteristics of response inhibition and perceptual decision-making, these two functions may closely be related. Specifically, response inhibition depends on perceived information of the external environment, and this information should be relevant to achieving response inhibition. Then, we asked how response inhibition and perceptual decision-making interact when task-relevant information involves perceptual uncertainty. In this context, neural mechanisms underlying response inhibition and perceptual decision-making may be coordinately involved in a brain-wide functional network. We hypothesized that there are three possibilities regarding the topological structure of the network: 1) a functionally merged region implements response inhibition under perceptual uncertainty (Shackman et al., 2011; Yarkoni et al., 2011); 2) distinct regions responsible for response inhibition (e.g., rIFC) and perceptual decision-making (e.g., MT) interact to guide response inhibition (Konishi et al., 1996; Egner and Hirsch, 2005; Kayser et al., 2010; Waskom et al., 2014; Tsumura et al., 2021a; Tsumura et al., 2021b); Fig 1B *middle*); and 3) a hub-like region links the regions involved in response inhibition and perceptual decision-making (Cole et al., 2013; Osada et al., 2015; Nee and D’Esposito, 2016; Jiang et al., 2018). To test these possibilities, functional MRI was performed while human participants performed a go/no-go task based on visually presented motion stimuli in which perceptual uncertainties was manipulated by the coherence of randomly moving dots (Figs. 1C/D). We first explored brain regions associated with response inhibition, motion strength, and their interaction. Then, we examined the directional effective connectivity between the task-related regions to examine their functional topological architecture.

## Materials and Methods

### Participants

Written informed consent was obtained from 26 healthy right-handed participants (10 females; age range: 18-22). Experimental procedures were approved by the institutional review boards of Keio University and Kochi University of Technology. Participants received 2000 yen for participation in each of two experimental sessions. Sample size was determined prior to the study based on behavioral piloting experiments.

### Session procedures

The experiment consisted of two sessions conducted on separate days. The first day involved practice sessions, where participants performed a discrimination task for motion dot stimuli (Figs. 1C/D). On the second day, they first performed the same discrimination task during fMRI scanning. Then, they performed a go/no-go task based on motion dot stimuli identical to those used in the discrimination task (Figs. 1C/D).

### Stimuli

All stimuli were generated with Matlab version 2012a, using the Psychophysics Toolbox extension, version 3.0.10 (Brainard, 1997; Pelli, 1997). The current stimuli were similar to those used in a previous study of perceptual decision-making (Chen et al., 2015) and task switching (Tsumura et al., 2021a; Tsumura et al., 2021b). Each motion stimulus was composed of 150 white dots moving inside a donut-shaped display patch, with a white cross in the center of the patch on a black background (Fig. 1C). The display patch was centered on the screen and extended from 6 to 12° of visual angle. Within the display patch, every dot moved at a speed of 10° of visual angle per second. Some dots moved coherently toward one direction while the others moved randomly. The percentage of coherently moving dots determined the “coherence”, which was set at one of three values (20, 40, and 80%). To remove local motion signals, dot presentation was controlled following a standard method for generating motion stimuli (Shadlen et al., 1996). Specifically, upon stimulus onset, new dots were presented at new random locations in each of the first three frames. Each dot was relocated after two subsequent frames, so that the dots in frame 1 were repositioned in frame 4, and the dots in frame 2 were repositioned in frame 5, etc. When repositioned, each dot was either randomly presented at the new location or aligned with the pre-determined motion direction (upward or downward), depending on the pre-determined motion strength in that trial. Each stimulus was composed of 18 video frames at 60 Hz refresh rates (i.e., 300-ms presentation). Before dot presentation, the color of the central fixation cross was changed to red from white in order to cue dot stimulus presentation. The cue stimulus was presented for 0.75 sec. The color of the fixation cross became white when dot stimuli disappeared and remained white for 1.75 sec.

### Experimental procedure

#### Direction discrimination task

At the beginning of each trial, a dot patch was presented, and participants were required to judge the direction of overall motion (up or down; Fig. 1D) and to press the corresponding button (left or right) with their right thumb as quickly and correctly as possible. The response window was 1050 ms. The stimulus-response relationship was identical the practice and scanning sessions, and was counterbalanced across participants.

Each participant completed six runs, and each run involved 70 trials. For familiarization, the first five trials in each run used the highest coherence (80%), and were not included in data analysis. The last five trials in each run also used high coherence and the results were discarded. Thus, in each run, the remaining middle 60 trials (20 trials for each coherence level) were analyzed.

#### Response inhibition task

Participants performed a go/no-go task (Fig. 1D) using random dot stimuli. As in the discrimination task, a dot patch was presented in each trial. Participants had to press a button as quickly as possible when the motion of the dot stimuli was upward (or downward) (go trial), whereas in trials with the alternative direction, they had to withhold the response (no-go trial). Motion directions for the go and no-go trials were counterbalanced across participants.

Each participant completed nine functional runs, each of which involved 70 trials. The first and last five trials of each run were high-coherence go trials, and their results were discarded. The remaining 60 trials involved 48 go trials and 12 no-go trials, and trials with each coherence level were equally distributed throughout one functional run (i.e. 20 trials for each coherence level).

### Imaging procedure

MRI scanning was performed by a 3-T MRI scanner (Siemens Verio, Germany) with a 32-channel head coil. Functional images were acquired using a multi-band acceleration gradient-echo echo-planar imaging sequence [repetition time (TR): 0.8 sec; echo time (TE): 30 msec; flip angle (FA): 45 deg; 80 slices; slice thickness: 2 mm; in-plane resolution: 2 x 2 mm; multiband factor: 8]. Each functional run involved 256 volume acquisitions. Data on the first 10 volumes were discarded to take into account the equilibrium of longitudinal magnetization. High-resolution anatomical images were acquired using an MP-RAGE T1-weighted sequence [TR: 2500 msec; TE = 4.32 msec; FA: 8 deg; 192 slices; slice thickness: 1 mm; in-plane resolution: 0.9 x 0.9 mm^2^].

### Behavioral analysis

Accuracy and reaction times were calculated for each of the two tasks (discrimination and go/no-go tasks) and each coherence level (20, 40, and 80%), and then compared across tasks and coherence levels. Statistical tests were performed based on repeated measures ANOVAs using SPSS Statistics 23 (IBM Corporation, NY USA).

To examine whether perceptual sensitivity to motion stimulus differed between the discrimination and go/no-go tasks, receiver operating characteristic (ROC) analysis (Luce et al., 1963) was performed. For each task and coherence level, differences in z-scores for hit and false alarm rates were calculated as d-prime values (Macmillian and Creelman, 1991).

### Image preprocessing

MRI data were analyzed using SPM12 software (http://fil.ion.ac.uk/spm/). All functional images were first temporally realigned across volumes and runs, and the anatomical image was coregistered to the mean of the functional images. The functional images were then spatially normalized to a standard MNI template with normalization parameters estimated for the anatomical scans. The images were resampled into 2-mm isotropic voxels, and spatially smoothed with a 6-mm full-width at half-maximum Gaussian kernel.

### General linear model (GLM) analysis

#### Single-level analysis

A GLM approach was applied to estimate parameter values for task events. The events of interest were correct no-go and go trials with the parametrical effect of coherence levels normalized (z-scored) across trials. For the go trials, normalized reaction times were also added as a parametrical effect. Error trials in the go and no-go trials were separately coded in GLM as nuisance effects. These task events were time-locked to the onset of motion dot stimuli and then convolved with a canonical hemodynamic response function implemented in SPM. Additionally, six-axis head movement parameters, white-matter signals, and cerebrospinal fluid signals were included in GLM as nuisance effects. Then, parameters were estimated for each voxel across the whole brain. To examine the effect of success in the no-go trials, a separate GLM analysis was performed in which the no-go trials were encoded with parametrical effects for success/error, coherence levels, and RTs (error trials only), normalized across trials.

For the discrimination task, a separate GLM analysis was performed. Event coding was similar to that in the go/no-go task, with correct trials coded separately based the direction used in the go/no-go task.

#### Group-level analysis

Maps of parameter estimates were first contrasted for each participant. The contrast maps were collected from all participants, and were subjected to group-mean tests based on non-parametric permutation methods (5000 permutations) implemented in *randomise* in the FSL suite (Eklund et al., 2016) (http://fmrib.ox.ac.uk/fsl/). Then voxel clusters were identified using a voxel-wise uncorrected threshold of P < .001. The voxel clusters were tested for significance with a threshold of P < 0.05 corrected by the family-wise error (FWE) rate. This procedure was empirically validated to appropriately control the false positive rate when correcting P-values across the whole brain (Eklund et al., 2016). The peaks of significant clusters were then identified and listed in tables. If multiple peaks were identified within 12 mm, the most significant peak was kept.

### Dynamic causal modeling (DCM) analysis

To examine task-related functional connectivity during the go/no-go task under perceptual uncertainty, DCM (Friston et al., 2003) analysis was performed. DCM allows us to explore effective connectivity among brain regions under the premise of the brain as a deterministic dynamic system that is subject to environmental inputs and that produces outputs based on a space-state model. DCM constructs a nonlinear system involving intrinsic connectivity, task-induced connectivity, and extrinsic inputs. Parameters of the nonlinear system are estimated based on fMRI signals (system states) and task events.

The MT is well known to be a neocortical temporal region that is specifically associated with motion perception (Shadlen et al., 1996; Beauchamp et al., 1997; Corbetta and Shulman, 2002; Huk et al., 2002). The rIFC is thought to be a core region involved in implementing response inhibition (Garavan et al., 1999; Konishi et al., 1999; Bokura et al., 2001; Braver et al., 2001; Menon et al., 2001; Aron et al., 2004; Aron and Poldrack, 2006; Li et al., 2006; Aron et al., 2007; Chikazoe et al., 2007; Li et al., 2008; Xue et al., 2008; Chambers et al., 2009; Chikazoe et al., 2009; Swann et al., 2009; White et al., 2014; Osada et al., 2019). The superior temporal sulcus (STS) showed prominent activity during high-coherent no-go trials in the current study (Fig. 4A and Table 3). Given existing evidence regarding the MT and rIFC and the current results for the STS, the regions of interest (ROIs) in the DCM analysis were defined as MT, STS, and rIFC.

These ROIs were created as 6 mm-radius spheres centered on peak coordinates determined by the following procedures that avoided circular analysis (Kriegeskorte et al., 2009). The center coordinate of the MT ROI was defined based on the group-level statistical map for the coherence effect during the discrimination task (Fig. 3A and Table 1), independently of our go/no-go data.

**Table 1.**
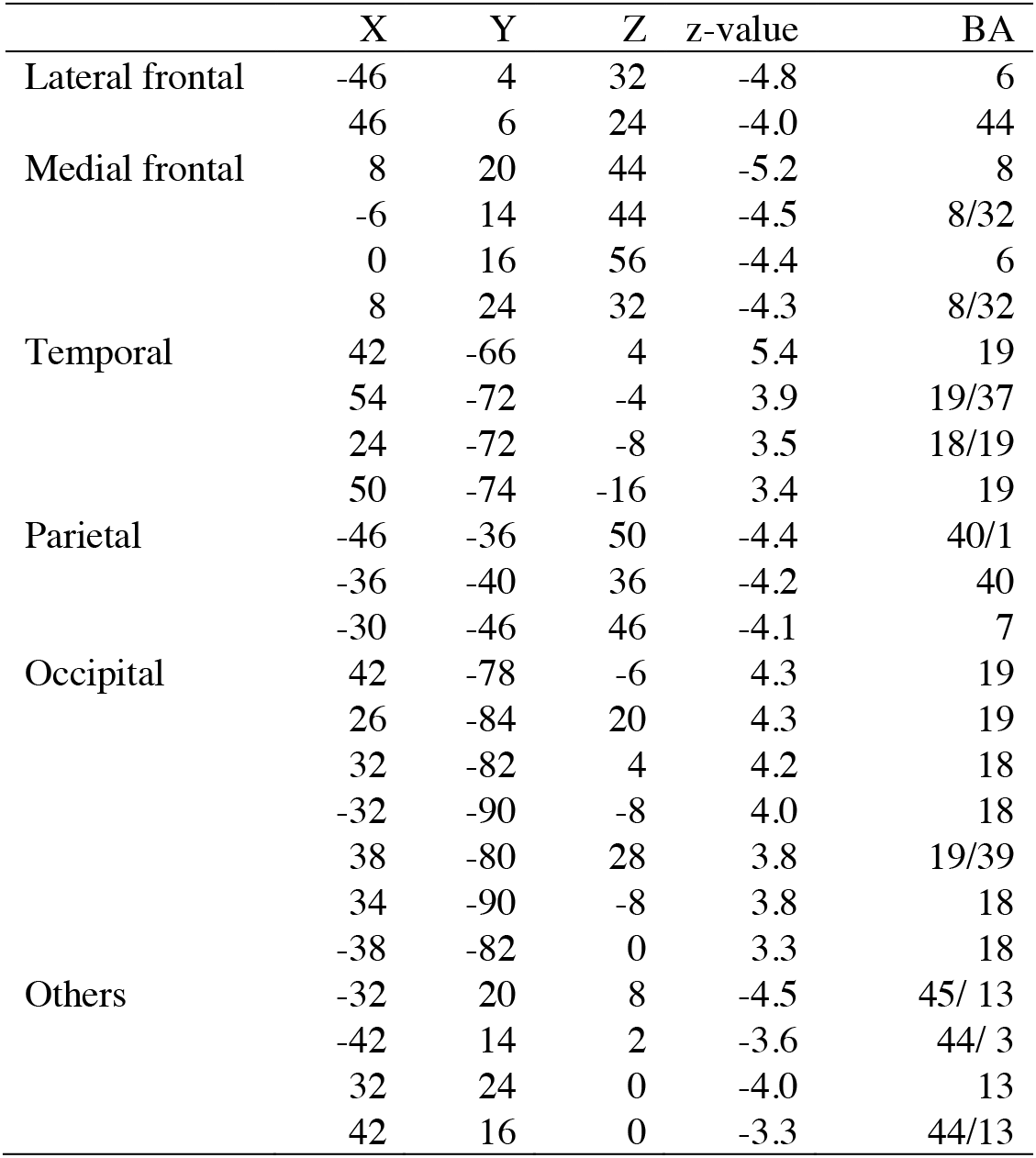
Brain regions showing a significant motion coherence effect in the discrimination task. Coordinates are listed in MNI space. Positive and negative z-values indicate an activity increase in high- and low-coherent trials, respectively. BA indicates Brodmann areas and is approximate.

The center coordinate of the rIFC ROI was defined based on a prior study of go/no-go (Zhang et al., 2017), independently of our go/no-go data. For reproducibility, another rIFC ROI was defined based on meta-analysis of response inhibition (Yarkoni et al., 2011) (http://www.neurosynth.org/analyses/terms/response%20inhibition/), and we confirmed that fundamental results were preserved.

For the STS, the center coordinate of the ROI was defined based on the coherence effect of the no-go trials relative to go trials using a leave-one-subject-out procedure. More specifically, the STS ROI was defined on an individual basis: for each participant, the coordinate was determined based on the group-level statistical map from which the participant was excluded when the group-level statistics were calculated (Tsumura et al., 2021a; Tsumura et al., 2021b).

The signal time courses of three ROIs and the regressors in events of interest were then extracted from first-level GLMs. Time courses are concatenated across functional runs, and nuisance effects of head-motion, white matter signals, ventricle signals, whole brain signals, functional runs, and contrast were subtracted out for ROI time courses.

For the no-go trials, causal models were defined as those that differed among ROIs in terms of external inputs and modulatory effects. Since the current analysis involved three ROIs, the tested models comprised 512 types (i.e., 2^3^ inputs and 2^6^ connection effects). Then, connectivity matrices reflecting 1) first-order connectivity, 2) effective changes in coupling induced by the inputs, and 3) extrinsic inputs to ROIs were estimated using SPM12 for each of 512 models based on DCM analysis. A parametric regressor for motion coherence level in the no-go trials was used as the extrinsic effect for effective connectivity between ROIs and ROI inputs.

To estimate the effective connectivity strength, a Bayesian model reduction method (Friston et al., 2016) was used. This reduction method enables the calculation of posterior densities of all possible reduced models, and these posterior densities were then inverted to a fully connected model. The reduced models were supplemented with the second-level parametric empirical Bayes method (Friston et al., 2016) to apply empirical priors that remove subject variability from each model.

Subsequently, parameters of these models were estimated based on Bayesian model averaging (Penny et al., 2010) to estimate connectivity strength. Because the current analysis aimed to identify the average effective connectivity observed across participants, we used a fixed effect (FFX) estimation assuming that every participant uses the same model, rather than a random effect (RFX) estimation assuming that different participants use different models; this latter estimation method has been often used to examine group differences in effective connectivity (Penny et al., 2010). The significance of connectivity was then tested by a posterior probability density and corrected for FWE rates across task-related connectivity patterns.

Additionally, to test the robustness of the functional connectivity, we examined BMA-based DCM without model reduction or an empirical prior, and confirmed that the overall results were consistent across these estimation methods (Tsumura et al., 2021a; Tsumura et al., 2021b).

## Results

### Behavioral results

In the discrimination task, accuracy was lower in the low-coherence (i.e., more uncertain) trials [F(1, 25) = 111.5; P < .001; Fig. 2A], replicating previous studies of perceptual decision-making (Shadlen et al., 1996; Kim and Shadlen, 1999; Mazurek et al., 2003; Palmer et al., 2005; Kayser et al., 2010).

**Figure 2.**
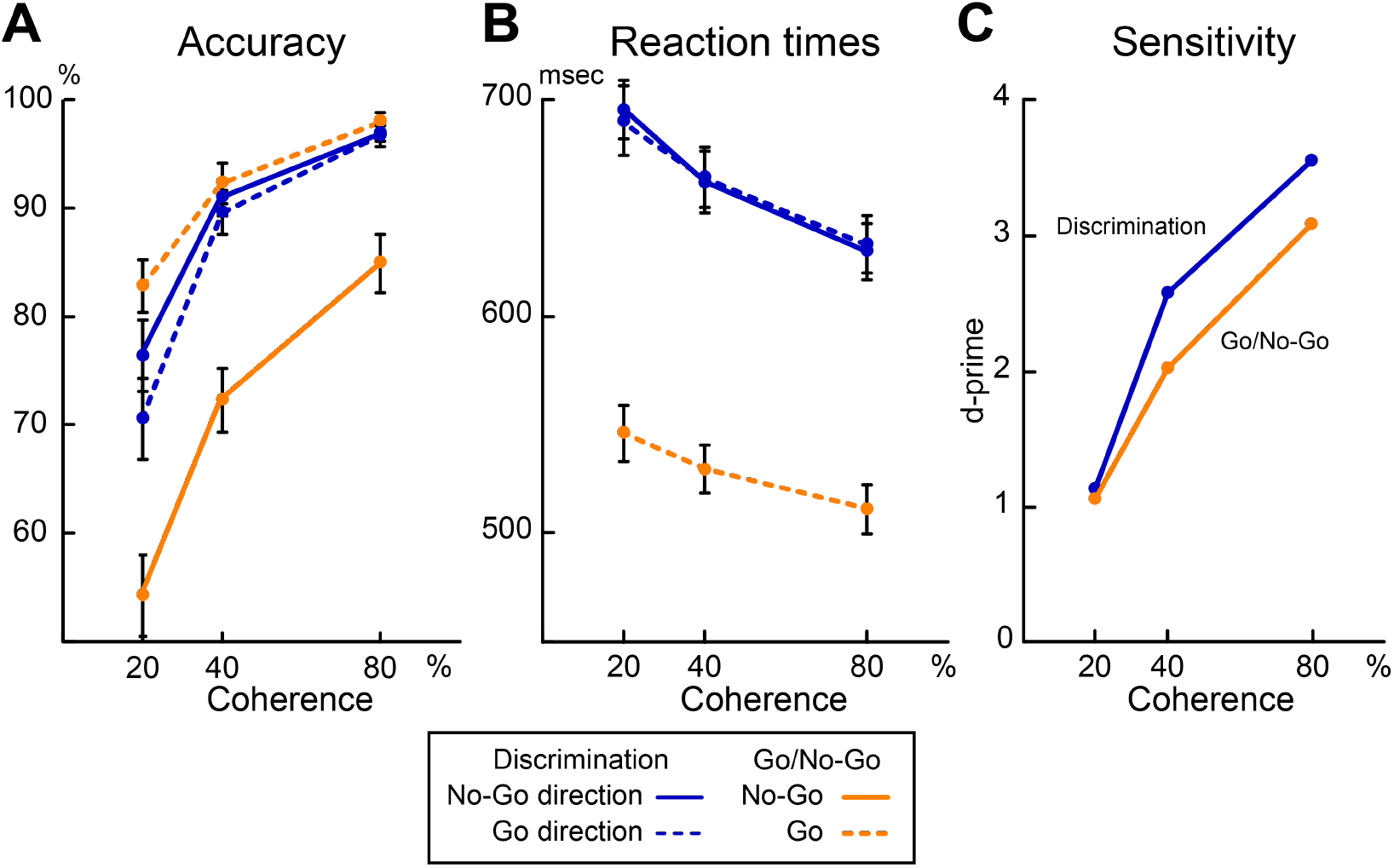
Behavioral results. A) Accuracy in the discrimination and go/no-go tasks as a function of motion coherence. The vertical and horizontal axes indicate accuracy and motion coherence, respectively. Blue and orange lines indicate the discrimination and go/no-go tasks, respectively. Solid and dotted lines indicate trials with no-go and go directions, respectively. B) Reaction times for the discrimination and go/no-go tasks. The vertical and horizontal axes indicate reaction times and motion coherence, respectively. C) Sensitivity to motion-dot stimuli during the discrimination and go/no-go tasks. The vertical and horizontal axes indicate d-prime values and motion coherence, respectively.

In the go/no-go task, accuracy was also lower in low coherence-trials [F(1, 25) = 146.6; P < .001; Fig. 2A]. In the no-go trials, accuracy was lower than in the go trials [F(1, 25) = 42.0; P < .001; Fig. 2A], consistent with previous studies of go/no-go tasks (Bokura et al., 2001; Chikazoe et al., 2009). Importantly, the accuracy in the no-go trials was lower than that in the direction-matched trials in the discrimination task [F(1, 25) = 62.7; P < .001; Fig. 2A].

To compare accuracy in the go trials and direction-matched trials in the discrimination task, a repeated measures ANOVA with the trial (go and direction-matched discrimination task trials) and the coherence levels (high, middle, and low) as factors was performed. Significant main effect of trials [F(1, 25) = 7.6; P < .05; Fig. 2A] and a significant interaction effect [F(1, 25) = 9.4; P < .01, planned contrast of a linear effect] were observed, indicating that in the go trials, accuracy was higher and coherence effect was weaker relative to those in the direction-matched discrimination task trials. This result suggests that prepotent tendency for go response was enhanced in the go/no-go task.

Then, we examined coherence effect in the no-go trial. A repeated measures ANOVA with the trial (go and no-go) and the coherence levels as factors revealed a significant interaction effect [F(1, 25) = 6.8; P < .05; linear coherence effect; Fig. 2A], indicating that the motion coherence effect was greater in the no-go trials, compared to the go trials. Likewise, a repeated measures ANOVA with the trial (no-go and direction-matched discrimination task trials) and coherence levels as factors revealed a significant interaction effect [F(1, 25) = 5.1; P < .05; linear coherence effect], again greater motion coherence effect in the no-go trials than in the discrimination task trials.

To test whether the greater coherence effect in the no-go trials is attributable to the change in perceptual sensitivity to motion stimuli or task demands, sensitivity to motion stimulus was examined based on d-prime indices for each task and coherence level (see Materials and Methods). In both the discrimination and go/no-go tasks, d-prime indices were greater in the high-coherence trials, consistent with the accuracy results (Fig. 2C). If the change in sensitivity to motion stimuli explains the greater coherence effect on accuracy in the no-go trials (i.e., lower accuracy in low coherence trials), the slope of d-prime along the coherence level would be steeper in the go/no-go task than in the discrimination task. However, the d-prime indices did not support this possibility because the d-prime slope was steeper in the discrimination task (Fig. 2B). These results suggest that the greater coherence effect in the no-go trials is attributable to differential task demands rather than the change in the sensitivity to motion between the go/no-go and discrimination tasks.

Reaction times (RTs) were shorter in high-coherence trials in both the discrimination task [F(1, 25) = 82.5; P < .001; linear coherence effect; Fig. 2B] and the go trials in the go/no-go task [F(1, 25) = 32.0; P < .001; linear coherence effect]. RTs were also shorter in the go trials than the trials in the discrimination task [F(1, 25) = 120.1; P < .001]. A repeated measures ANOVA with trials (go trials and direction-matched discrimination trials) and coherence levels as factors revealed a significant interaction effect [F(1, 25) = 6.9; P < .05; linear coherence effect].

These collective behavioral results suggest that the current behavioral task successfully manipulated response inhibition and perceptual decision-making, and more importantly, the motion coherence affected both perceptual decision-making and response inhibition.

### Imaging results

We first explored brain regions associated with motion coherence. As shown in Fig. 3A and Table 1, prominent activation was observed in the middle temporal (MT) cortex in high-coherence trials, consistent with prior studies (Shadlen et al., 1996; Beauchamp et al., 1997; Corbetta and Shulman, 2002; Huk et al., 2002; Tsumura et al., 2021a; Tsumura et al., 2021b). On the other hand, in low-coherence trials, activation was greater in multiple frontal regions including inferior frontal cortex (IFC) and pre-supplementary motor area (pre-SMA), also consistent with previous studies (Zatorre et al., 1994; Gevins et al., 1997; Duncan and Owen, 2000; Paulus et al., 2001; Lavie, 2005; Vickery and Jiang, 2009; Graves et al., 2010; Sheth et al., 2012; Tsumura et al., 2021a; Tsumura et al., 2021b).

**Figure 3.**
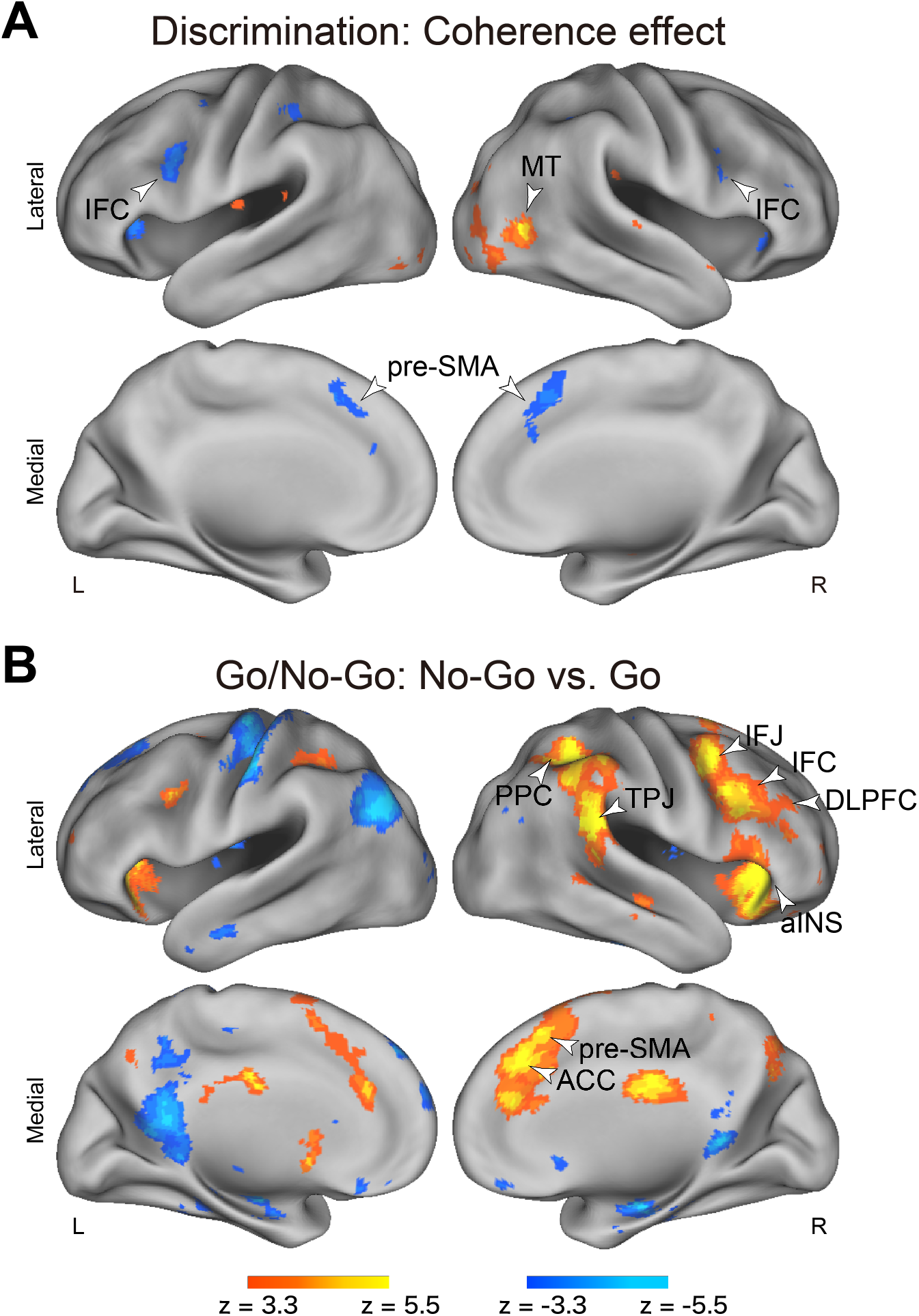
A) Statistical maps of the parametrical effect of motion coherence in the discrimination task. Maps are overlaid onto a 3D surface of the brain. Hot and cool colors indicate positive and negative effects, respectively. The white arrows indicate the MT (middle temporal), IFC (inferior frontal cortex), and pre-SMA (supplementary motor area) regions. B) Statistical activation maps for signal increases and decreases in the no-go trials relative to go trials. Hot and cool colors indicate signal increases and decreases, respectively, in the no-go trials. Formats are similar to those in Figure 3A. The white arrowheads indicate the IFC, MT, pre-SMA, aINS (anterior insular), DLPFC (dorsolateral prefrontal cortex), IFJ (inferior frontal junction), PPC (posterior parietal cortex), TPJ (temporoparietal junction), and ACC (anterior cingulate cortex) regions.

We next explored brain regions associated with response inhibition. In the no-go trials, relative to the go trials, robust activity was observed in fronto-parietal regions including the IFC, inferior frontal junction (IFJ), dorsolateral prefrontal cortex (DLPFC), anterior insula (aINS), anterior cingulate cortex (ACC), pre-supplementary motor area (pre-SMA), posterior parietal cortex (PPC) and temporo-parietal junction (TPJ), predominantly in the right hemisphere (Fig. 3B and Table 2). These results are consistent with prior studies of response inhibition (Garavan et al., 1999; Rubia et al., 2003; Aron et al., 2004; Li et al., 2006; Chambers et al., 2009; Chikazoe et al., 2009). Importantly, the rIFC cluster overlapped with a rIFC cluster showing significant activation during lower coherent trials in the discrimination task (147 voxels), suggesting that the rIFC is associated with both response inhibition and greater decision demands in low-coherence trials, consistent with a previous study (Tsumura et al., 2021a).

**Table 2.**
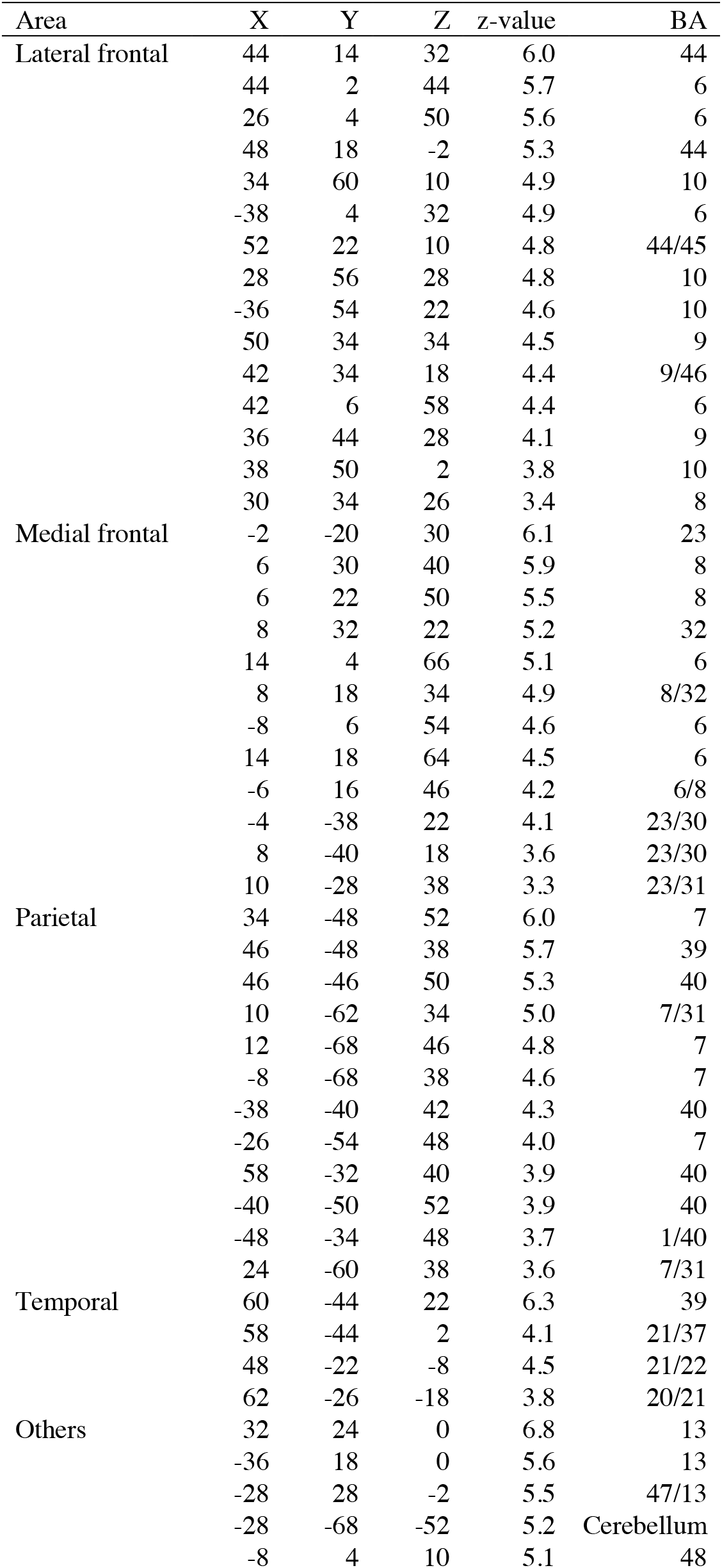
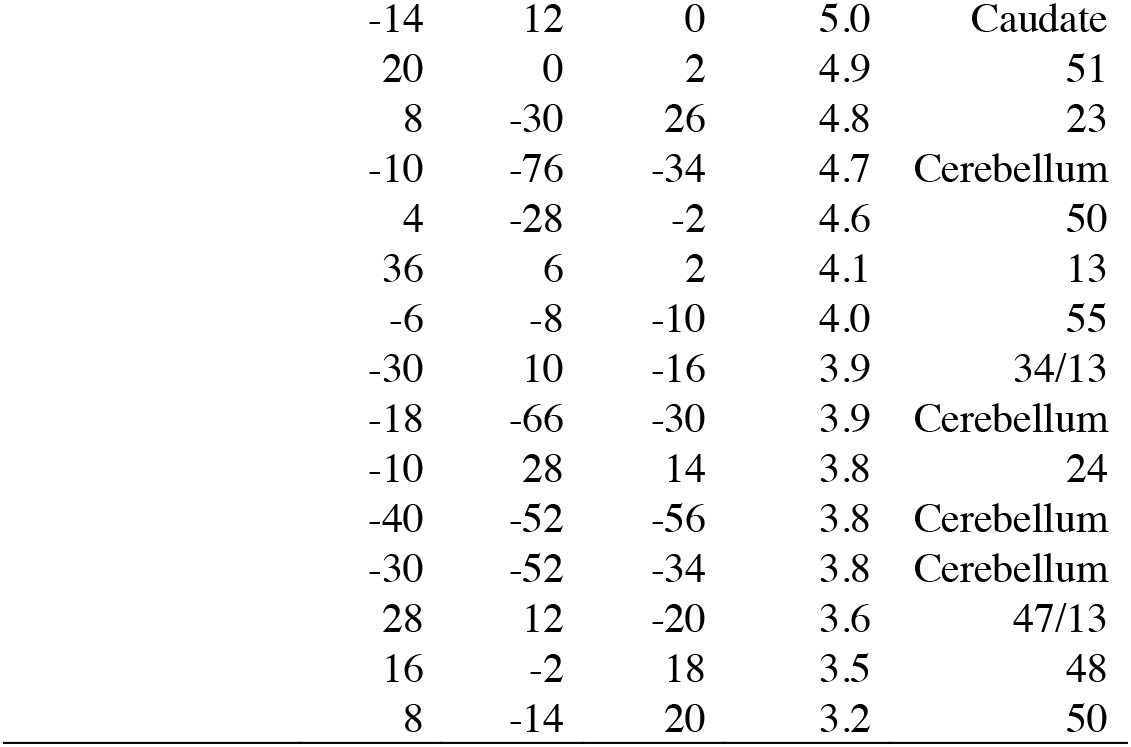
Brain regions showing a significant signal increase in the no-go trials relative to the go trials in the go/no-go task. Formats are similar to those in Table 1.

Then, we explored brain regions showing significant interaction effect of motion coherence and response inhibition during the go/no-go task by contrasting parametric modulatory effect of motion coherence between the no-go and go trials (Fig. 4A and Table 3). A robust difference in coherence effect was observed in the right superior temporal sulcus (STS). Notably, this area showed greater activation in the high-coherence no-go trials relative to low-coherence no-go trials (Fig. 4B *left* and Table 4) and in the no-go relative to go trials (Fig. 3B/4B *right* and Table 2), collectively indicating that the STS activity was enhanced specifically during high-coherence no-go trials.

**Figure 4.**
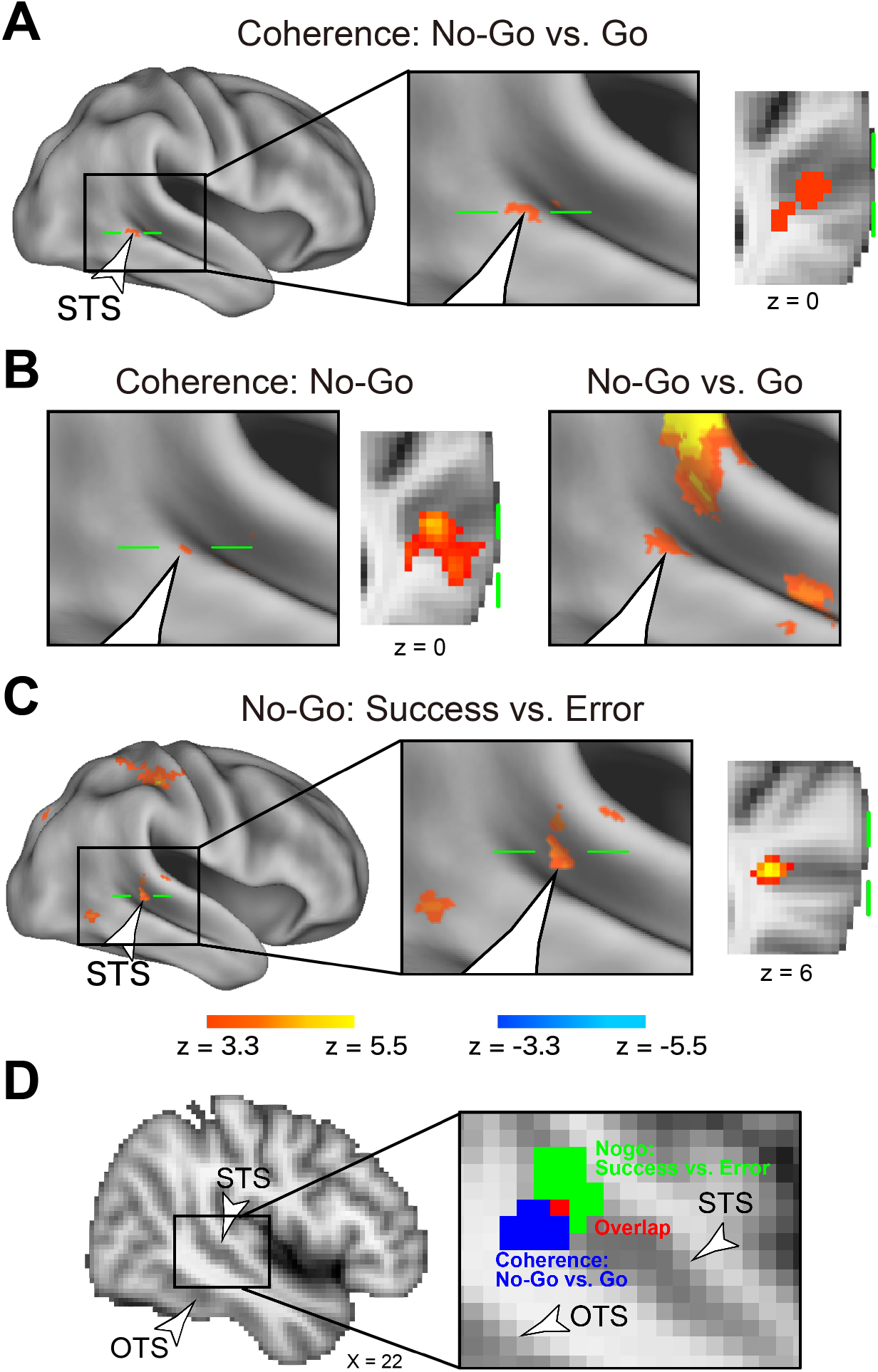
A) Statistical maps for differential parametrical effects of motion coherence between the no-go vs. go trials. Hot and cool colors indicate greater positive coherence effects in the no-go and go trials, respectively. The area enclosed in the black rectangle (*left*) represents the area magnified on the middle. The green solid line on the surface indicates the axial section on the right. Formats are similar to those in Figure 3A. The white arrowheads indicate the STS region. B) Statistical maps for parametrical effects of motion coherence for the no-go trials (*left*) and for activation during the no-go trials relative to go trials (*right*; Fig. 3B). Formats are similar to those in Figure 3A *bottom*. C) Statistical activation map for signal increases and decreases indicating success and error in the no-go trials. Formats are similar to those in Figure 3A. The white arrowheads indicate the STS region. D) The STS region shows greater activation in successful no-go trials (green), and coherence effects in the no-go trials are mapped onto a 2D parasagittal slice of the standard brain. The section is indicated by the X coordinate on the bottom right. OTS: occipitotemporal sulcus.

**Table 3.**
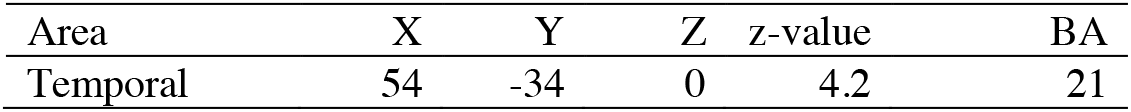
Brain regions showing a significant difference in the coherence effect between the no-go and go trials. Positive z-values indicate a greater coherence effect in the no-go trials. Formats are similar to those in Table 1.

**Table 4.**
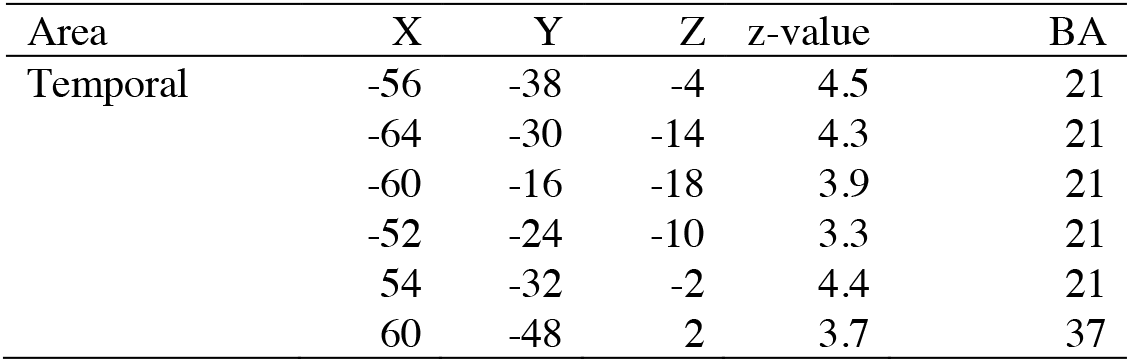
Brain regions showing a significant coherence effect in the no-go trials. Positive and negative z-values indicate an increase in high- and low-coherence trials, respectively. Formats are similar to those in Table 1.

| Area | X | Y | Z | z-value | BA |
| --- | --- | --- | --- | --- | --- |
| Temporal | -56 | -38 | -4 | 4.5 | 21 |
|  | -64 | -30 | -14 | 4.3 | 21 |
|  | -60 | -16 | -18 | 3.9 | 21 |
|  | -52 | -24 | -10 | 3.3 | 21 |
|  | 54 | -32 | -2 | 4.4 | 21 |
|  | 60 | -48 | 2 | 3.7 | 37 |

The STS involvement in high-coherence no-go trials may point to an important role of this region in both response inhibition and perceptual decision-making. In particular, STS activity during response inhibition is elevated in situations where perceptual decision-making is more facilitated, suggesting that the STS is specifically affected by perceptual decision-making during response inhibition. We then hypothesized that the success of response inhibition depends at least in part on the STS playing a hub role between perceptual decision making and response inhibition. To test this hypothesis, we compared brain activity between success and error trials in the no-go trials, and whole-brain exploratory analysis revealed robust activation in the right STS (Fig. 4C and Table 5). Of note, this STS region is located very close to the STS region that showed a strong coherence effect in the no-go trials (Fig. 4D), with the two areas only 12.2-mm apart (Tables 2/5). Taken together, our results highlight important roles of the STS in both response inhibition and perceptual decision-making.

**Table 5.** Brain regions showing a significant activity increase in the correct no-go trials relative to the error no-go trials. Positive and negative z-values indicate an activity increase in the correct and error trials, respectively. Formats are similar to those in Table 1.

| Area | X | Y | Z | z value | BA |
| --- | --- | --- | --- | --- | --- |
| Temporal | 54 | -36 | 12 | 4.8 | 22 |
|  | 64 | -28 | 12 | 3.9 | 22/41 |
| Parietal | 38 | -30 | 46 | 4.9 | 1 |
|  | -2 | -38 | 66 | 4.5 | 5 |
|  | 26 | -46 | 50 | 4.3 | 7 |
|  | 32 | -46 | 62 | 4.2 | 1/5 |
|  | 44 | -20 | 56 | 4.0 | 1/4 |

### Directional effective connectivity

The whole-brain exploratory analyses identified three types of brain regions: 1) the MT, associated with perceptual decision-making (Fig. 3A); 2) the rIFC, associated with response inhibition (Fig. 3B); and 3) the STS, associated with both of them (Figs. 4A/B). In order to examine the functional relationships between these regions during response inhibition under perceptual uncertainty, we performed interregional effective connectivity analysis of coherence effects in the no-go trials based on dynamic causal modelling (Friston et al., 2019). DCM allowed us to examine the directionality of task-related functional connectivity based on state-space models.

Figure 5 shows the task-related parametrical modulatory effect of motion coherence in the no-go trials. In the high coherence no-go trials, connectivity was enhanced from the MT toward the rIFC via the STS (Ps < .05, FWE corrected). In contrast, connectivity was enhanced from the rIFC to the MT and STS in low-coherence no-go trials (Ps < .05, FWE corrected; Fig. 2). These results suggest a reversal of the connectivity between rIFC, STS, and MT regions depending on motion coherence: bottom-up signaling from the MT to the rIFC via the STS under low perceptual uncertainty, which reversed to top-down signaling from the rIFC to the STS and MT regions under high perceptual uncertainty.

**Figure 5.**
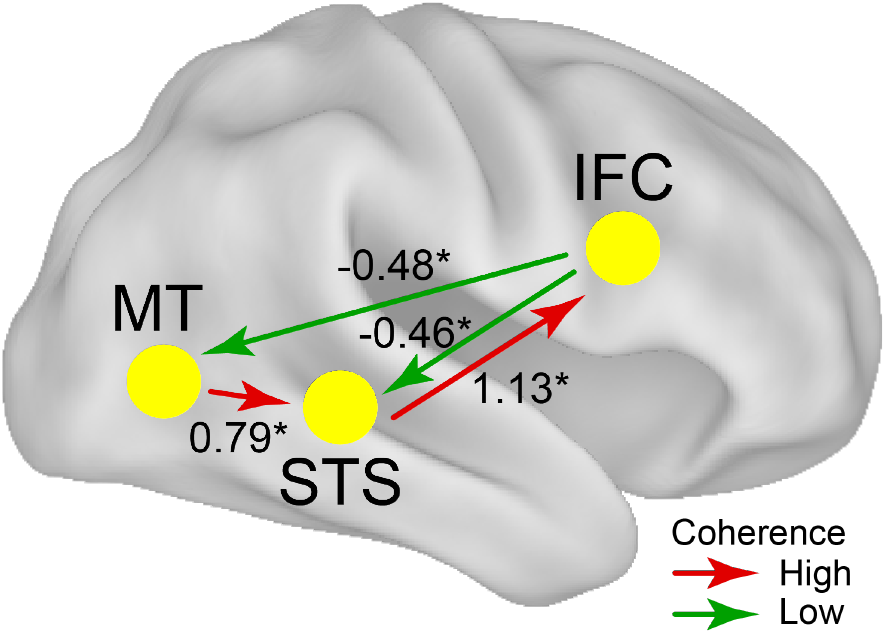
Effective connectivity analysis. Red and blue arrows indicate enhancements of effective connectivity in high- and low-coherence trials, respectively, as reflected by positive and negative values of the estimated connectivity strength. *: P < .05, FWE corrected.

## Discussion

The current study provides new insights regarding the relationships between perceptual decision-making and response inhibition. Our results highlighted the important role of the STS in high-coherence no-go trials and in the success of response inhibition (Fig. 6), and support Hypothesis 3 (Fig 1A; see Introduction). The STS serves as a hub-like region that links the rIFC, which is involved in response inhibition, and the MT, which play a role in perceptual decision-making. Specifically, in high-coherence trials, the STS received task-related signals from the MT that was associated with motion perception, and signaled the rIFC regarding response inhibition. The signal direction was reversed in low-coherence trials, such that the rIFC signaled the MT and STS. These results demonstrate that bottom-up and top-down signals are reversible, and their status depends on the uncertainty of perceptual information (Fig. 6). As such, the fronto-temporal network may convert perceptual information to executive control information via the STS to achieve response inhibition.

**Figure 6.**
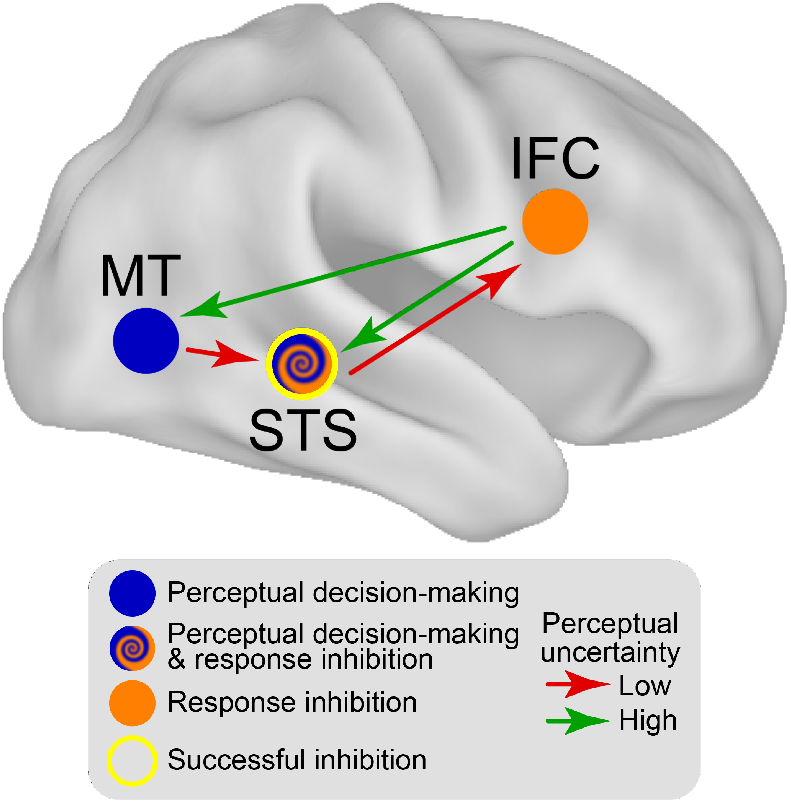
Schematic diagrams of neural mechanisms among the STS, IFC and MT regions during response inhibition with perceptual uncertainty. Arrows indicate signal directions.

### The STS links perceptual and executive information

It is known that the MT receives motion-related information from early visual cortices and is more activated when motion coherence is higher (Desimone and Ungerleider, 1989; Ungerleider and Haxby, 1994; Desimone and Duncan, 1995; Tootell et al., 1995; Ungerleider et al., 1998). It is also known that the rIFC and pre-SMA are activated during response inhibition and signaled to basal ganglia (Rubia et al., 2003; Duann et al., 2009; Cai and Leung, 2011; Jahfari et al., 2011). In the current study, the STS was activated during successful high-coherence no-go trials; it may receive perceptual information from the MT, and send signals to the rIFC, where response inhibition is encoded. Thus, the STS may encode both perceptual decision-making and response inhibition and serve as a hub role between them (Fig. 6).

Prior studies have suggested that top-down and bottom-up signaling between neocortical temporal regions and prefrontal regions are implicated in executive control (Kawamura and Naito, 1984; Wilson et al., 1993; Desimone and Duncan, 1995; Tomita et al., 1999; Moore and Armstrong, 2003). Our task-related effective connectivity analysis demonstrated that which of these two types of signaling was active depended on perceptual uncertainty, as demonstrated in our recent study (Tsumura et al., 2021b). Additionally, the STS relayed signals from the MT to rIFC, suggesting a functional route from the temporal cortex to the frontal cortex.

It is well-known that visual information coded in the temporal cortex becomes more abstract along the posterior-to-anterior axis of the cortical areas (Desimone and Ungerleider, 1989; Ungerleider and Haxby, 1994; Desimone and Duncan, 1995; Ungerleider et al., 1998). For example, when a face stimulus is perceived, posterior temporal regions represent visual features of the face, whereas anterior regions represent the identity of the face (Freiwald and Tsao, 2010). This hierarchical information stream in the temporal cortex is consistent with the current findings. In particular, the MT, which is located in the posterior end of the inferior sulcus, represented visual perceptual information (Huk et al., 2002), and the STS, which is located anteriorly, represented a mixture of visual and control information that was converted into executive control information in the rIFC.

Many studies of visual motion perception have consistently reported the involvement of the MT (Newsome and Paré, 1988; Britten et al., 1992, 1993; Zohary et al., 1994; Shadlen et al., 1996; Beauchamp et al., 1997; Britten and Newsome, 1998; Kim and Shadlen, 1999; Braddick et al., 2001; Corbetta and Shulman, 2002; Huk et al., 2002; Mazurek et al., 2003), but some neuroimaging studies have also reported the STS is active during the perception of visual motion stimuli (Braddick et al., 2001; Noguchi et al., 2005; Kayser et al., 2010; Tsumura et al., 2021a; Tsumura et al., 2021b). This STS involvement may partially reflect executive control processing, as is shown in the present study. Indeed, previous studies of response inhibition have reported involvement of the lateral temporal areas, including the temporo-parietal junction (Aron and Poldrack, 2006; Xue et al., 2008; Chikazoe et al., 2009) and distributed regions along the superior temporal sulcus (Aron et al., 2007; Chikazoe et al., 2009). It is possible that these temporal regions convert visual information into executive information along the bottom-up stream in the neocortical temporal areas (Osada et al., 2021).

### Perceptual decision-making and executive control

Our recent studies examined the relations between perceptual decision-making and task switching, another representative executive control function (Tsumura et al., 2021a; Tsumura et al., 2021b). In these studies, to examine task switching under perceptual uncertainty, we manipulated the motion coherence of similar random dot stimuli, as in the current study, while participants alternated discrimination tasks.

In one study (Tsumura et al., 2021a), the left lateral prefrontal cortex was implicated in task switching, and stimulus-modality-dependent occipitotemporal regions received complementary signals from the right lateral prefrontal cortex involved in perceptual decision-making, supporting Hypothesis 2 in Fig. 1A. In another study (Tsumura et al., 2021b), top-down signaling from the prefrontal cortex supplemented task-relevant information in the occipitotemporal regions, also supporting Hypothesis 2.

In contrast, the current results support Hypothesis 3. One helpful way to explain the variability in the possible topological architecture of the functional network is to examine whether the executive control functioning is affected by the perceptual decision-making. This can be statistically tested by interaction effects of motion coherence and executive control. Behaviorally, interaction effects between motion coherence and task switching were absent in our previous studies, indicating that the task-switching effect itself was not affected by the motion coherence level. Consistent with the behavioral results, neuroimaging analysis did not reveal significant interaction effects of task switching and motion coherence.

In the current study, by contrast, accuracy showed an interaction effect between response inhibition (no-go vs. go) and motion coherence, with decreased accuracy in lower coherence trials, particularly the no-go trials (Fig. 2A). The STS region showed a consistent interaction effect, with greater activity in high-coherence no-go trials. These behavioral and neural interaction effects between perceptual decision-making and response inhibition may reflect the hub role of the STS. As such, the STS may relay task-related information from the MT to the rIFC by converting perceptual information in the MT into response inhibition information in the rIFC, thus successfully exerting executive control to achieve a behavioral goal.

### Prefrontal involvements in high decision demands

In low-coherence trials, the MT was less activated than in high-coherence trials, reflecting the slower accumulation of perceptual information (Newsome and Paré, 1988; Britten et al., 1992, 1993; Zohary et al., 1994; Shadlen et al., 1996; Beauchamp et al., 1997; Britten and Newsome, 1998; Kim and Shadlen, 1999; Braddick et al., 2001; Corbetta and Shulman, 2002; Huk et al., 2002; Mazurek et al., 2003; Kayser et al., 2010). On the other hand, fronto-parietal regions such as the IFC, IFJ, and pre-SMA, and PPC were activated in low coherence trials in the current discrimination task (Fig. 3A) and in our previous studies (Tsumura et al., 2021a; Tsumura et al., 2021b). This strong prefrontal involvement may reflect greater decision demands for low-coherence motion stimuli, which is consistent with increased frontal activation in high cognitive load situations (Zatorre et al., 1994; Gevins et al., 1997; Duncan and Owen, 2000; Paulus et al., 2001; Lavie, 2005; Vickery and Jiang, 2009; Enriquez-Geppert et al., 2010; Graves et al., 2010; Sheth et al., 2012). Our results suggest that such strong prefrontal involvement in low-coherence trials supplements the perceptual information in the MT, consistent with our previous studies (Tsumura et al., 2021a; Tsumura et al., 2021b).

## Acknowledgements

This study was supported by JSPS Kakenhi, 26350986, 26120711, 17H05957, and 17K01989 to KJ, 20H00521 and 21K18267 to MT, 17H00891 to KN, and 17H06033 to JC, a grant from Uehara Memorial Foundation to KJ, and a grant from Takeda Science Foundation to KJ. We thank Drs. Russell A. Poldrack, Alexander C. Huk, and Corey N. White for scientific advice. We also thank Maoko Yamanaka for administrative assistance.

